# *Metacollinia emscheri* n. sp., a novel sanguicolous apostome ciliate of freshwater amphipods (*Gammarus* spp.)

**DOI:** 10.1101/2024.07.10.602951

**Authors:** Sebastian Prati, Sonja Rückert, Daniel S. Grabner, Bernd Sures, Jamie Bojko

## Abstract

We describe a novel sanguicolous parasitic ciliate, *Metacollinia emscheri* n. sp., found in the freshwater amphipods *Gammarus pulex* and *G. fossarum*. This ciliate infected up to 20% of the amphipods collected in a German stream catchment, the Boye, a tributary of the river Emscher. The ciliate showed morphological characteristics fitting the genus *Metacollinia*. Different life stages of variable size occurred simultaneously in the hemocoel throughout the hosts’ body. The tomont had 40-47 slightly spiraled kineties, a non-ciliated cortical band, a large macronucleus, and contractile vacuoles arranged in rows or scattered throughout the cytoplasm. The protomites/tomites with nine somatic kineties presented evidence of the buccal kineties x, y, and z reminiscent of those of the order Foettingeriida. Phylogenetic analyses of the 18S rRNA and COI regions confirm the ciliate placement in the Collinidae and a close relatedness to the type species of the genus *Metacollinia*, *Metacollinia lucensis*.

We formally describe this new parasite as *Metacollinia emscheri* n. sp. using pathological, morphological, and nuclear/mitochondrial genetic data. The systemic infections observed in histological preparations and the pathogenicity of *Metacollinia emscheri* n. sp. suggest that this parasite might influence host population dynamics. Given the ecological importance of amphipods as keystone species in freshwater ecosystems, an outbreak of this parasite might indirectly impact ecosystem functioning.

## 1. Introduction

Macroinvertebrates, including amphipods, are essential components of stream ecosystem food webs. Amphipods are highly diversified and ubiquitous keystone organisms that often represent the dominant element of invertebrate communities (Giari et al., 2020; Macneil et al., 1997). Their shredding activity can account for up to 75% of overall leaf-litter breakdown in some ecosystems, and they are an important food source for a wide range of predators, linking several trophic levels within riverine food webs (Giari et al., 2020). Their ecological niche makes them ideal hosts for a hyper-diverse range of pathogens ranging from viruses to trophically transmitted metazoans (Bojko and Ovcharenko, 2019; Warren et al., 2023, 2022). Yet most studies focus on metazoan parasites, gregarines (McKinley et al. 2024), or microsporidians due to their abundance, diversity, and ecological relevance (Prati et al., 2023).

Protists, such as ciliates, are commonly found as commensal epibionts on the carapace, gills, and extremities of amphipods, where they benefit from the host movement to collect particulate matter (Bojko et al., 2013). However, few apostome ciliates have specialized symbiotic and parasitic relationships with crustaceans, including amphipods (Lynn, 2008). Among them, sanguicolous parasites are able to penetrate the host cuticle and feed on the cells and fluid of the hemocoel (Lynn et al., 2014; Lynn and Strüder-Kypke, 2019). Representatives of the Collinidae and the Pseudocollinidae families reproduce continuously in the host hemolymph, leading to systemic infections, which might ultimately result in host death (Chantangsi et al., 2013; Lynn et al., 2014; Lynn and Strüder-Kypke, 2019; Prokopowicz et al., 2010). For instance, Pseudocollinidae representatives have been linked to mass mortality events in krill in marine environments (Gómez-Gutiérrez et al., 2012; Gómez-Gutiérrez and Kawaguchi, 2017) and the Collinidae member *Lynnia grapsolytica* strongly enhances the mortality rate of infected crabs under laboratory conditions (Metz and Hechinger, 2021).

The majority of Collinidae are parasites of freshwater amphipods and isopods (De Puytorac and Lom, 1962). One of the first observations in freshwater amphipods dates back to 1852 when an apostome ciliate, later described as *Paracollinia branchiarum* was found in the hemolymph of *Gammarus pulex* (De Puytorac and Lom, 1962). Successive investigations revealed the presence of Collinidae in freshwater amphipods from Lake Baikal and subterranean environments in Europe and the USA (Capriulo and Small, 1985). More recently, an unidentified ciliate has been histologically observed in the hemolymph of *Gammarus roeselii* (Bojko et al., 2017). These findings indicate that these parasites might be ubiquitous but poorly investigated.

Given the ecological importance of amphipods and the possible high pathogenicity of ciliates belonging to the Collinidae family, deepening our knowledge of these sanguicolous parasites is paramount. If highly pathogenic, these parasites might strongly influence host population dynamics and indirectly affect ecosystem functioning. We document the pathology, morphological characteristics, and genetic detail of a novel member of the Collinidae, in the hemocoel of the amphipods *Gammarus pulex* and *Gammarus fossarum,* and formally describe it as *Metacollinia emscheri* n. sp.

## 2. Materials and methods

### 2.1 Sampling

Amphipods were collected between December 2022 and March 2024 from eight sampling locations across the Boye catchment in North Rhine-Westphalia, Germany (Table 1). The first collection of amphipods (December 2022) initially aimed at describing novel microsporidian parasites comprised 72 individuals (n = 52 from BOYohKI and n = 20 from VORohBOY, Table 1). Thus, only microsporidian-infected amphipods (n = 20; for molecular identification method, see Prati et al. 2023) were processed for histological analyses. After finding an individual infected with an unknown sanguicolous ciliate, we decided to carry out a non-targeted sampling in April 2023.

**Table 1.** Sampling locations, sampling date, amphipod species, average size of the 4^th^ coaxal plate in mm with SD, and prevalence in % of *Metacollinia emscheri* n. sp.

| Location | Coordinates | Date | N | Species | Size in mm | Prevalence in % |
| --- | --- | --- | --- | --- | --- | --- |
| BOYohBR | 51.569135; 6.927698 | 27/04/2023 | 4 | *Gammarus pulex* clade C | 2.88 ± 0.21 | 0 |
|  |  |  | 26 | *Gammarus pulex* clade E | 2.95 ± 0.46 | 0 |
| BOYohKI | 51.553107; 6.948059 | 07/12/2022 | 8 | *Gammarus pulex* clade C | 2.57 ± 0.31 | 12.50 (1/8) |
|  |  |  | 4 | *Gammarus pulex* clade E | 2.49 ± 0.37 | 0 |
|  |  | 13/05/2023 | 18 | *Gammarus pulex* clade C | 2.63 ± 0.43 | 0 |
|  |  |  | 12 | *Gammarus pulex* clade E | 2.71 ± 0.41 | 0 |
| BOYohSP | 51.564297; 6.930277 | 27/04/2023 | 10 | *Gammarus fossarum* clade 2 | 2.82 ± 0.56 | 0 |
|  |  |  | 8 | *Gammarus pulex* clade C | 2.99 ± 0.74 | 0 |
|  |  |  | 12 | *Gammarus pulex* clade E | 2.96 ± 0.57 | 0 |
| BOYuhHA | 51.534515; 6.997740 | 02/05/2023 | 26 | *Gammarus fossarum* clade 2 | 3.08 ± 0.51 | 11.53 (3/26) |
|  |  |  | 4 | *Gammarus pulex* clade C | 3.28 ± 0.66 | 0 |
| BRA1000 | 51.577238; 6.932752 | 27/04/2023 | 1 | *Gammarus pulex* clade C | 3.36 | 0 |
|  |  |  | 25 | *Gammarus pulex* clade E | 2.95 ± 0.38 | 0 |
| HAAob | 51.570306; 6.960891 | 13/05/2023 | 11 | *Gammarus pulex* clade C | 2.87 ± 0.30 | 9.09 (1/11) |
|  |  |  | 19 | *Gammarus pulex* clade E | 2.81 ± 0.36 | 5.26 (1/19) |
| HAAun | 51.562638; 6.955581 | 13/05/2023 | 9 | *Gammarus pulex* clade C | 3.02 ± 0.31 | 22.22 (2/9) |
|  |  |  | 21 | *Gammarus pulex* clade E | 2.92 ± 0.49 | 19.05 (4/21) |
|  |  | 02/03/2024 | 42 | Not determined | 3.07 ± 0.47 | 21.43 (9/42) |
| VORohBOY | 51.554753; 6.932435 | 07/12/2022 | 8 | *Gammarus pulex* clade C | 2.84 ± 0.29 | 0 |
|  |  | 13/05/2023 | 25 | *Gammarus pulex* clade C | 3.15 ± 0.63 | 0 |
|  |  |  | 5 | *Gammarus pulex* clade E | 3.07 ± 0.71 | 0 |

All amphipods were kept alive and dissected within 6h post collection. The head and anterior part of the thorax were fixed in 96% ethanol for molecular analyses. The proximate segments of the thorax (2 mm) were placed in 2.5% glutaraldehyde 0.1 M sodium cacodylate buffer (pH 7.4) for electron microscopy. The rest of the body was fixed in Davidsońs freshwater fixative for 24 hours, then transferred to 70% ethanol for histological examination. Additionally, given the high morphological variability observed in the ciliates, we collected 42 amphipods in March 2024. We dissected them and separately stored the freshly isolated ciliates by life stages in 96% ethanol for molecular analyses to confirm their taxonomic placement.

### 2.2 Histological analyses

Amphipod tissues were infiltrated with paraffin wax in an automated tissue processor (ethanol-xylene substitute-wax), embedded into wax blocks, and sectioned to 4 µm thickness through the animal center. Each section was mounted on a glass slide and stained with hematoxylin and eosin, following a standard de-wax, stain, and rehydration protocol. The slides were screened for pathogens using an Olympus CX 33 trinocular light microscope and photographed with a Sony Alpha 6600 mounted via extension tubes on the camera port.

### 2.3 Scanning Electron Microscopy

Haemocoel from glutaraldehyde-fixed or fresh amphipods was transferred into self-made filter baskets with a 10 μm polycarbonate membrane filter (Millipore Corp.) attached to a cut-off BEEM® conical tip capsule. These capsules were placed in 0.9% NaCl in a multi-well plate. A piece of filter paper was mounted on the inside of the multi-well plate lid and saturated with 4% (w/v) OsO_4_. The ciliates were fixed with OsO_4_ vapor for 30 min in the dark. Ten drops of 4% (w/v) OsO_4_ were added directly to the filter baskets, and the samples were fixed again for 30 min. The filters with the ciliates were washed with distilled water, dehydrated with a graded series of ethyl alcohol, and critical point dried with CO_2_ (Leica EM CPD 300 critical point drier). Filters were mounted on stubs, sputter coated with 5 nm platinum (Leica ACE 600 sputter coater), and viewed using the Zeiss Crossbeam 540 scanning electron microscope.

### 2.4 Quantitative protargol staining

We employed a rapid Quantitative protargol staining (QPS) on ciliates isolated from glutaraldehyde-fixed amphipods to better observe the internal structures of ciliates. The protocol (see details in Supplementary material 1) is a modified version of those proposed by Skibbe (1994), Kurilov (2017), and Ji and Wang (2018). Briefly, the steps employed involved in-house production of liquid protargol (Kurilov, 2017), the concentration of fixed ciliates on cellulose filters (Skibbe, 1994), coverage of ciliates with a thin layer of agarose, hardening of agarose with formaldehyde, fast impregnation in warm protargol solution (Skibbe, 1994), dehydration and mounting of the filter on permanent slides. The process included steps to correct overbleaching and overstaining (Ji and Wang, 2018).

### 2.5 Molecular analyses

DNA was isolated from muscular tissue following a modified salt precipitation protocol described by Grabner et al. (2015). DNA extracted from the only amphipod infected with an unknown ciliate in December 2022 was sequenced using shotgun next-generation sequencing (NGS) to assemble the ciliate mitogenome. Based on the obtained sequences, we then conducted a molecular screening of sanguicolous ciliates in amphipods collected in April 2023 using DNA barcoding (Sanger sequencing). DNA barcoding was also used to confirm that all ciliate life stages isolated in March 2024 belonged to the same species. Furthermore, we molecularly identified infected hosts using DNA barcodes.

### 2.6 Genomic library preparation, sequencing, and bioinformatics

A DNA sample from an animal with histological confirmation of the infection was used for shotgun next-generation sequencing (NGS). The DNA extract was submitted to Novogene, where a library was prepared (NEBNext® Ultra™ DNA Library Preparation Kit), which results in paired-end (150 bp) sequence data. The library was sequenced on an Illumina NovaSeq. Data were returned in fastq format (raw forward reads: 34.6 M; raw reverse reads: 34.6 M) and trimmed and quality checked using Trimmomatic (Bolger et al., 2014). Assembly of the data was conducted in SPAdes v.3.15.3 (Bankevich et al., 2012), resulting in a contiguous sequence file containing 107,638 contigs (>1000 bp) and assembly statistics of: N50: 830; N90: 545; L50: 181,716; L90: 460,369; determined using quast v.5.2.0 (Gurevich et al., 2013).

The contigs were screened for mitochondrial sequences using a combination of metaxa2 v.2.1.3 (Bengtsson-Palme et al., 2015) and blastp (based on a bespoke protein database developed from available Ciliophora mitogenomes). This isolated two contigs that contributed to the genome of the suspected ciliate observed in histological section. These contigs were mapped to confirm contiguity (using CLC v.12) and annotated using Mfannot (Lang et al., 2023; translation table 4 - mold and protozoan).

### 2.7 DNA barcoding

For the molecular screening of sanguicolous ciliates in amphipods collected in April 2023 and for the ciliates isolated in March 2024, we used the primer pair: SB2F (5- TGTAAAACGACGGCCAGTGTTCCCCTTGAACGAGGAATTC-3) and ITS2.2R (5-CAGGAAACAGCTATGACCCTGGTTAGTTTCTTTTCCTCCGC-3) in conjunction with the pair cil18SspannerF (5-TGTAAAACGACGGCCAGT-AATCAGAGTAATGATTAATAGG-3) and cil18SspannerR (5-CAGGAAACAGCTATGAC-TGGTTCACCGGACCACTCGA-3); which targets the 18S rRNA and LSU rRNA region (Adlard and Lester, 1995; Metz and Hechinger, 2021) as these primers bound well to the mitogenome sequences obtained from the ciliate collected in December 2022 and did not amplify amphipods DNA. All infected amphipods were identified using the universal eukaryotic primers LCO1490 and HCO2198, which amplify the COI region.

Each PCR reaction volume used for the SB2F-ITS2.2R and cil18SspannerF- cil18SspannerR primer pairs consisted of 20 μL composed of 10 μL of 2 × AccuStart II PCR ToughMix (Quantabio), 1 μL of each primer (0.5 μM), 0.35 μL of 50 × GelTrack Loading Dye (Quantabio), 6.65 μL MilliQ water and 1 μL of DNA template. For the LCO1490 and HCO2198 primer pair, the PCR reaction consisted of 20 μL assay with 10 μL of OneTaq Quick-Load 2x Master mix (New England Biolabs), 1 μL of each primer (0.5 μM), 7 μL of nuclease-free water and 1 μL of DNA template. The PCR settings for the SB2F- ITS2.2R and cil18SspannerF- cil18SspannerR primers were as follows: initial denaturation at 94°C for 5 min, 35 cycles of denaturation at 94°C for 45 sec, annealing at 55°C for 75 sec, and extension at 72°C for 90 sec, with a final extension at 72°C for 7 min. PCR setting for LCO1490 and HCO2198 primers were as follows: initial denaturation at 94 °C for 60 sec, five cycles of denaturation at 94 °C for 60 sec, annealing at 45 °C for 90 sec, and extension at 68 °C for 90 sec, followed by 40 cycles at 94 °C for 60 sec, 50 °C for 90 sec, 68 °C for 60 sec, with a final extension at 72 °C for 5 min. PCR products were sent to Microsynth Seqlab (Germany) for sequencing using the forward primers SB2F, cil18SspannerF, and LCO1490.

### 2.8 Genetic distance and phylogenetic analyses

Raw 18S rRNA sequences (infected amphipods collected in April 2023 and ciliate isolated in March 2024) and a COI sequence (extracted from the mitogenome assembly) were quality-checked and edited using Geneious Prime v2024.0.5 (Biomatters). The obtained sequences were separately aligned with apostome ciliate sequences available on GenBank (www.ncbi.nlm.nih.gov) using the MAFFT v7.490 algorithm with standard settings (Katoh et al., 2019). All available 18S rRNA sequences with a minimum length of 1500 bp were used for genetic distance analyses, whereas 18S rRNA and COI sequences with a minimum length of 750 bp were used for phylogenetic analyses. Genetic distance was calculated in Mega11 v11.0.13 (Tamura et al., 2021) from 1450 unambiguously aligned nucleotide positions using the Kimura two-parameter (K2P) model with 1000 bootstrap replicates. Maximum likelihood phylogenetic trees with bootstrap support values (1000 replicates) were produced in IQ-Tree 2.2.2.6 (Minh et al., 2020). The HKY+F+I+R2 (18s) and TPM2u+F+I+G4 (COI) substitution models were selected based on Bayesian information criterion scores. The ciliates *Cinetochilum ovale* (Genbank accession number FJ870103) and *Paramecium bursaria* (JX082062) were used respectively as outgroups for 18S rRNA and COI sequences. Ciliates and their hosts were identified by comparing obtained sequences against GenBank records using Blastn.

## 3. Results

A total of 256 amphipods collected between December 2022 and April 2023 from eight sampling sites were screened for sanguicolous ciliates molecularly and histologically (Table 1). All known amphipod species occurring in the Boye system (Prati et al., 2022) were represented in the sample. More specifically, *G. fossarum* clade 2 (n = 36, 100% similarity, 100% coverage, e-value = 0 to ON093817), *G. pulex* clade C (n = 96, 99.9-100% similarity, 100% coverage, e-value = 0 to ON093813), and clade E (n = 124, 99.9-100% similarity, 100% coverage, e-value = 0 to ON093815). The most common amphipods in the investigated sites were *G. pulex* clade E and C.

### 3.1 Prevalence

Half of the sampling sites had ciliate-infected amphipod populations, with overall prevalences ranging from 6.67 to 21.43% (Table 1). Among the 12 infected amphipods collected between December 2022 and April 2023, three were *G. fossarum* clade 2, four *G. pulex* clade C, and 5 *G. pulex* clade E. Two ciliate-infected individuals presented simultaneous infections with Microsporidium sp. 515. One of them was also infected with a trematode. The 9 infected amphipods collected in March 2024 and used to isolate fresh ciliates were neither molecularly identified nor screened for co-infections. Overall, the average size of infected amphipods (4^th^ coaxal plate length 3.07 mm ± 0.44 SD, n = 21) was similar to that of uninfected ones (2.93 mm ± 0.49 SD, n = 277). No individual below 2.57 mm was infected.

### 3.2 Histology and pathology

Histological analysis indicated that the ciliate is a parasite that travels through the hemolymph and causes a systemic infection. Freely dividing tomonts and protomites-tomites were visible in the host hemocoel, surrounding the testis, muscle, hepatopancreas, and gill lamellae (Fig. 1). We found no evidence that the ciliate infiltrated these organs or any sign of visible immune reactions in the form of melanization. However, compared to uninfected individuals, the muscular mass progressively reduced with increasingly high ciliate densities in infected individuals.

**Fig 1.**
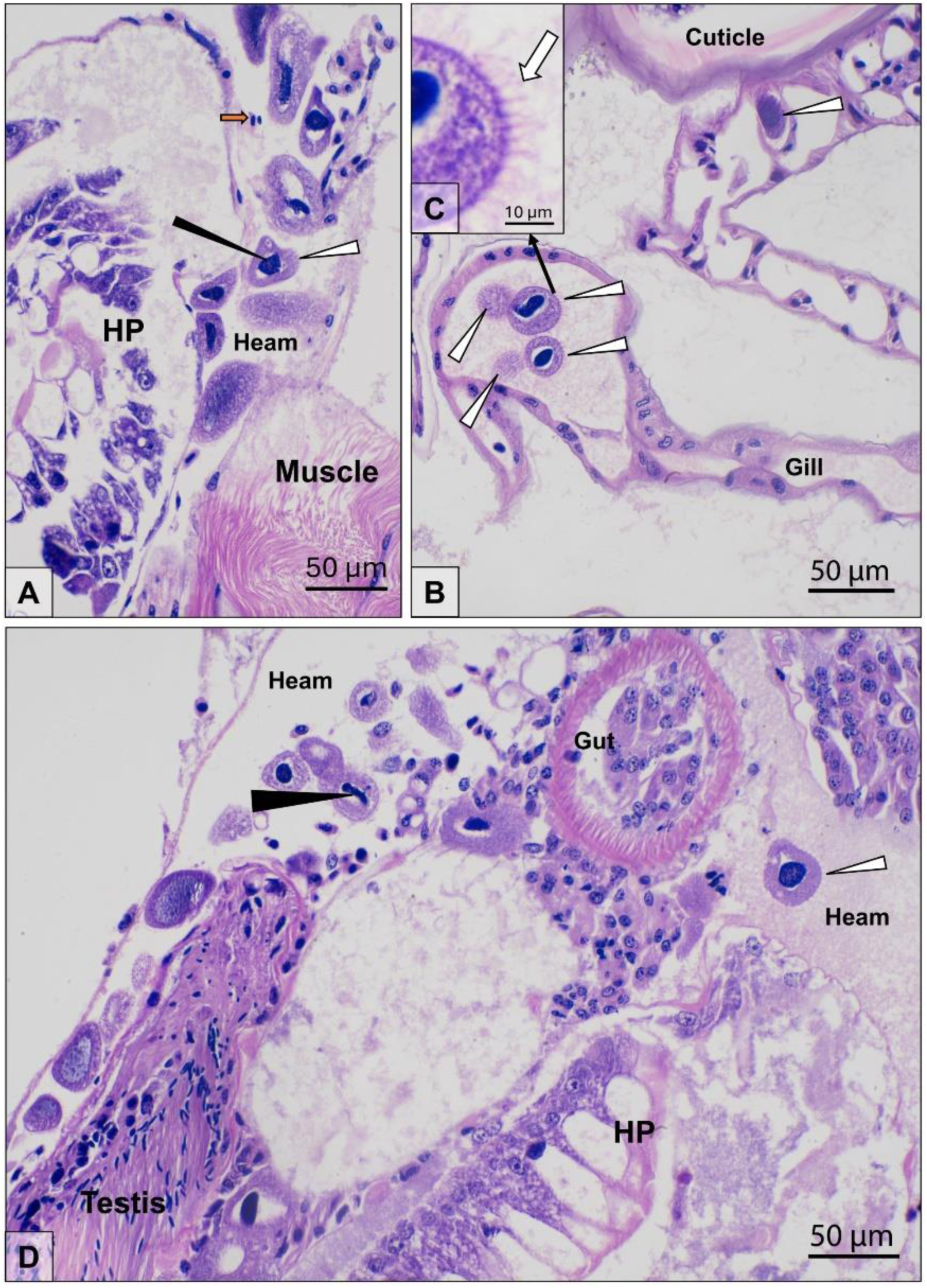
Histopathological preparations of the ciliate infection in amphipod tissue. A) Ciliated protozoans (white triangle) with basophilic staining nuclei (black triangle) are present in the hemocoel of the infected amphipod, alongside hemocytes (orange arrow). The image includes hemocoel (’Heam’) surrounding the hepatopancreas (’HP’) and muscle. B) Ciliated protozoans (white triangle) passing through the gill lamellae (’Gill’), alongside the outer host cuticle. C) Ciliate-like features are observable (white arrow) on the outside of the protozoans. D) Ciliated protozoans (white triangle) seen to travel through the hemocoel (’Heam’) adjacent to the testis, gut, and hepatopancreas (’HP’). An individual in section highlights the common U-shaped nucleus often noted in Ciliophora members.

### 3.3 Morphology

Similar to the histological sections, we observed tomonts and protomites-tomites in the glutaraldehyde-fixed gammarids. The tomonts were subelliptical and had a dense ciliature. Their length ranged between 34.6 and 90.8 µm (66.8 µm ± 15.3 SD), and the width between 15.4 and 59.1 µm (32.5 µm ± 8 SD). Tomonts had a large central macronucleus (width 48.5 µm ± 13.2 SD and length 15.9 µm ± 4.8 SD) and contractile vacuoles either arranged in rows parallel or slightly diagonal to kineties or rather scattered throughout the cytoplasm (Fig. 2 and 3). The largest tomonts displayed 40 to 47 irregularly spaced kineties converging toward the extremities but ending shortly before reaching the poles (Fig. 2 and 3). However, the vast majority of tomonts were along the path of regressive division; thus, it is possible that non-regressing cells might attain a larger size and kineties number than what is reported here.

**Fig 2.**
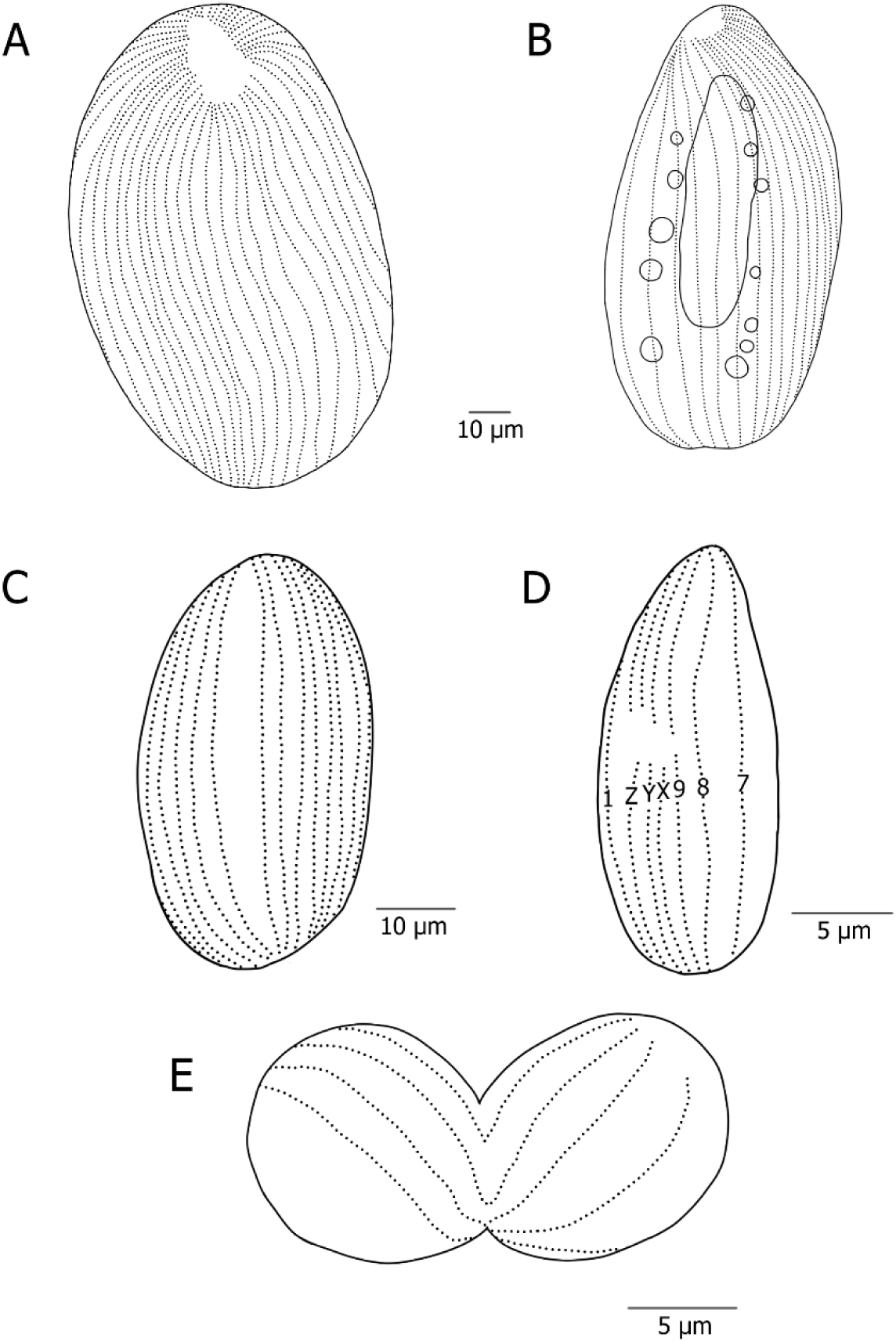
A) tomont with 47 kineties, B) internal morphology of tomont showing macronucleus and contractile vacuoles, C) regressing tomont displaying a non-ciliated cortical band, D) dividing protomite showing 9 kineties and the x, y, and z kineties, and E) conjugant.

**Fig 3.**
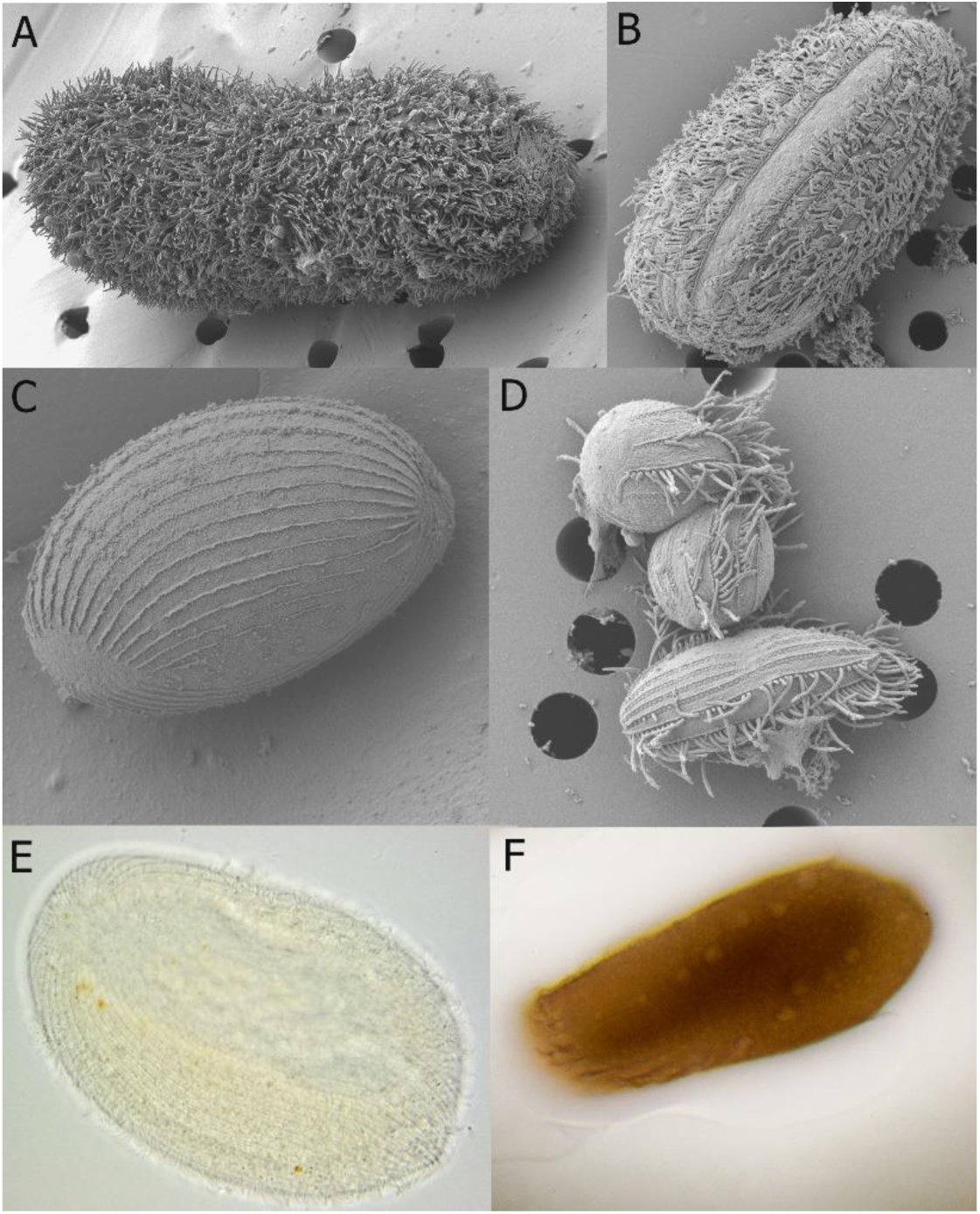
A) dividing tomont, B) regressing tomont displaying a non-ciliated cortical band, C) tomont showing kineties merging toward the poles but not touching them, D) dividing tomite and conjugant, E) light microscopy of freshly isolated tomont with kineties and a large macronucleous visible, and F) QPS staining of tomont showing contractile vacuoles arranged in parallel lines and a large macronucleous.

Following palintomy, a process during which the parental cell freely undergoes a sequence of repeated divisions without intervening growth in the host hemocoel, the size and the number of kineties of the daughter cells are progressively reduced. Regressing tomonts showed a non-ciliated band likely resulting from the reabsorption of kineties (Fig. 2 and 3). The protomites-tomites shape varied from pyriform to subelliptical, and their size ranged between 9.9 and 27.8 µm (20.7 ± 4.6) in length and 9.3 to 23 µm (14 ± 2.9) in width. They had a large macronucleus (length 11.1 ± 3.1 µm and width 6.73 ± 1.8 µm), nine somatic kineties, and in some cases, presented tree buccal kineties indicated as x, y, and z (Fig. 2 and 3).

### 3.4 Molecular phylogeny and genetics

Molecular identification of the collected sanguicolous ciliates using aligned 18S rRNA sequences indicated that these were closely related to *Metacollinia luciensis* (MH200618, 95.90% similarity, 100% coverage, e-value = 0.0), and all belonged to the same haplotype (Fig. 4). Similar results were obtained for the COI region (MH182621, 75.50% similarity, 99% coverage, e-value = 1e- 140); however, the availability of sequences for this region was very limited (Fig. 4). Molecular identification of all ciliate life stages isolated from amphipods collected in March 2024 confirmed that these belonged to the same species and haplotype. The genetic divergence (K2P) between the sequences obtained in the current study (0 differing nucleotides) and those of the closest related species *M. lucensis* (69 SNPs) and *Lynnia grapsolytica* (74 SNPs), both belonging to the Collinidae family, were respectively 4% and 4.7%, suggesting a closer genetic relationship with *M. luciensis*. Apostome ciliates belonging to other families diverged by 6.4-6.7% (92-100 SNPs), supporting the placement of the ciliate in the Collinidae.

**Fig 4.**
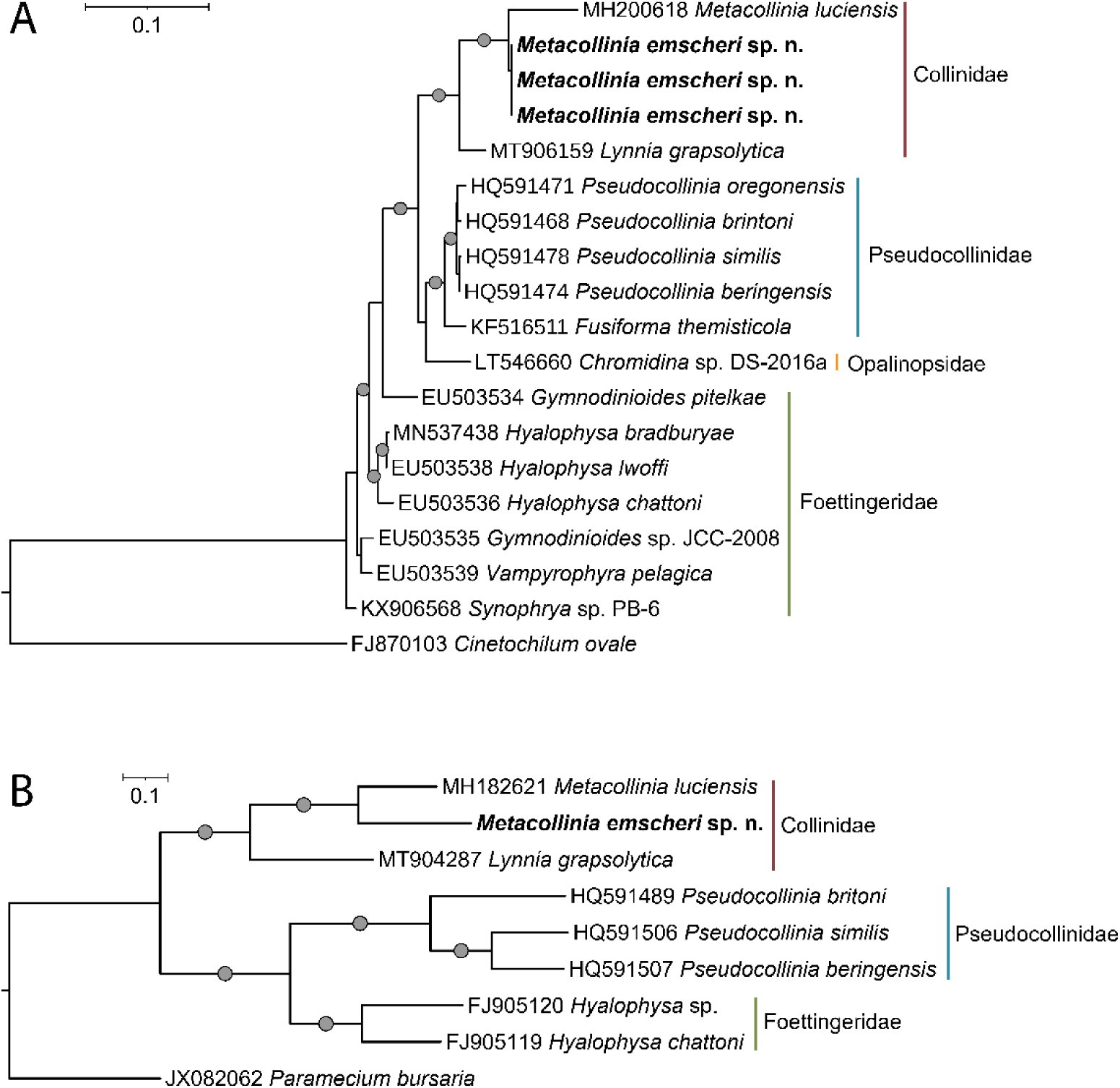
A) maximum-likelihood phylogeny of Apostome ciliates incorporating 18S rDNA and B) mitochondrial COI sequences. The tree includes the ciliates *Cinetochilum ovale* (18s) and *Paramecium bursaria* (COI) as outgroup. Best-fit model: HKY+F+I+R2 (18s) and TPM2u+F+I+G4 (COI) chosen according to Bayesian Information Criterion. Gray dots indicate bootstrap support values above 90 (1,000 replicates).

### 3.5 Mitochondrial genetics

The mitochondrial genome of the new ciliated protozoan was identified in two separate contigs, the larger consisting of 39,710 bp and encoding 28 protein-coding genes (PCGs); and the second was composed of 20,221 bp and encoded 4 PCGs, as well as both the SSU and LSU rRNA sequences. tRNA sequences remain unannotated and require more detailed exploration since predictive methods were unable to clearly identify putative coding regions. The most similar known relatives are *Metacollina*, *Tetrahymena*, and *Cryptocaryon* where proteins are compared from both contigs using blastp (Table 2).

**Table 2.**
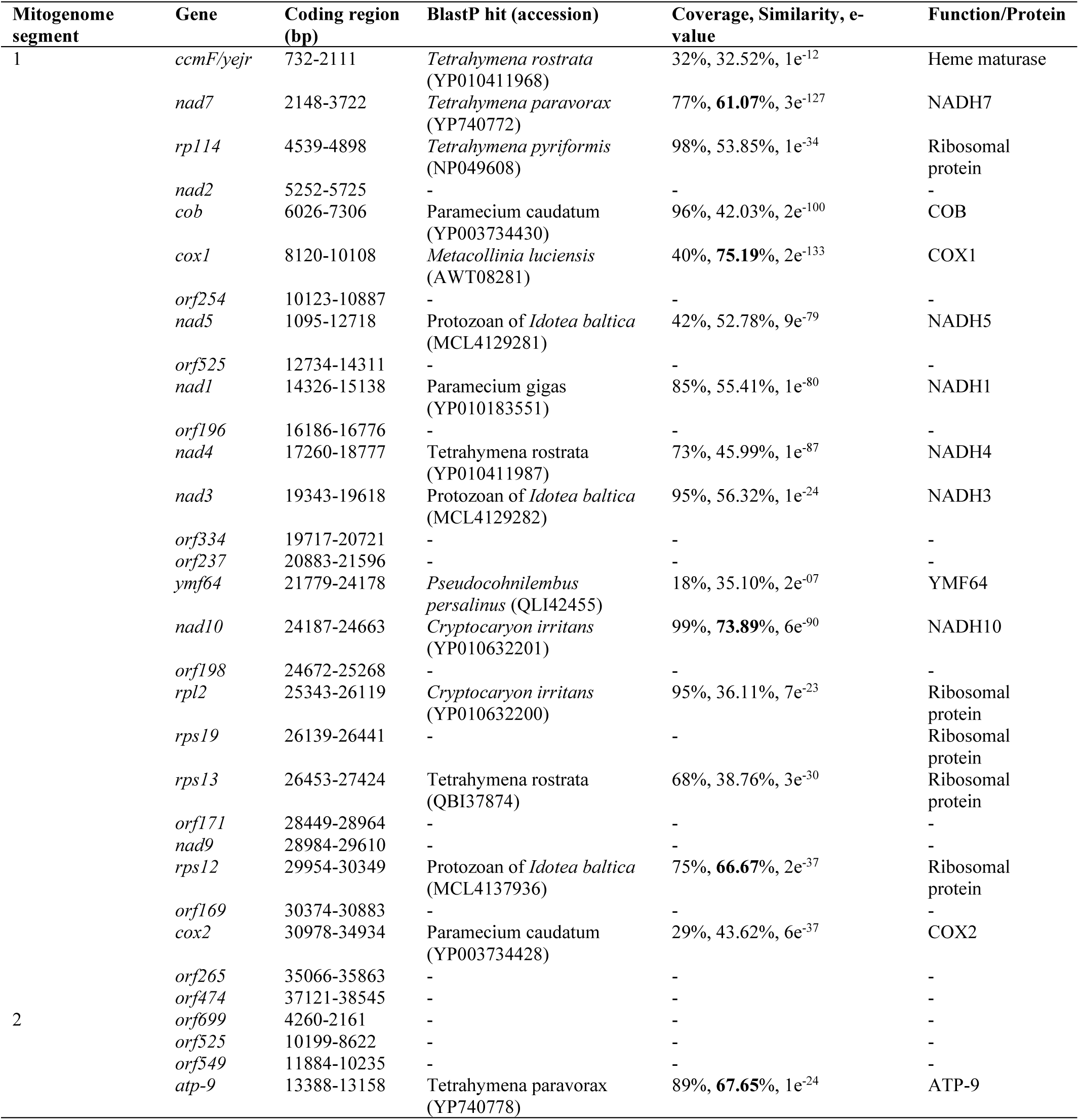
A blast table comparing the protein coding genes (as determined by MFannot) from both segments of the mitochondrial genome to available sequences held by NCBI.

| Mitogenome segment | Gene | Coding region (bp) | BlastP hit (accession) | Coverage, Similarity, e-value | Function/Protein |
| --- | --- | --- | --- | --- | --- |
| 1 | *ccmF/yejr* | 732-2111 | *Tetrahymena rostrata* (YP010411968) | 32%, 32.52%, 1e<sup>-12</sup> | Heme maturase |
|  | *nad7* | 2148-3722 | *Tetrahymena paravorax* (YP740772) | 77%, **61.07%**, 3e<sup>-127</sup> | NADH7 |
|  | *rpl14* | 4539-4898 | *Tetrahymena pyriformis* (NP049608) | 98%, 53.85%, 1e<sup>-34</sup> | Ribosomal protein |
|  | *nad2* | 5252-5725 | - | - | - |
|  | *cob* | 6026-7306 | *Paramecium caudatum* (YP003734430) | 96%, 42.03%, 2e<sup>-100</sup> | COB |
|  | *cox1* | 8120-10108 | *Metacollinia luciensis* (AWT08281) | 40%, **75.19%**, 2e<sup>-133</sup> | COX1 |
|  | *orf254* | 10123-10887 | - | - | - |
|  | *nad5* | 1095-12718 | Protozoan of *Idotea baltica* (MCL4129281) | 42%, 52.78%, 9e<sup>-79</sup> | NADH5 |
|  | *orf525* | 12734-14311 | - | - | - |
|  | *nad1* | 14326-15138 | *Paramecium gigas* (YP010183551) | 85%, 55.41%, 1e<sup>-80</sup> | NADH1 |
|  | *orf196* | 16186-16776 | - | - | - |
|  | *nad4* | 17260-18777 | *Tetrahymena rostrata* (YP010411987) | 73%, 45.99%, 1e<sup>-87</sup> | NADH4 |
|  | *nad3* | 19343-19618 | Protozoan of *Idotea baltica* (MCL4129282) | 95%, 56.32%, 1e<sup>-24</sup> | NADH3 |
|  | *orf334* | 19717-20721 | - | - | - |
|  | *orf237* | 20883-21596 | - | - | - |
|  | *ymf64* | 21779-24178 | *Pseudocohnilembus persalinus* (QLI42455) | 18%, 35.10%, 2e<sup>-07</sup> | YMF64 |
|  | *nad10* | 24187-24663 | *Cryptocaryon irritans* (YP010632201) | 99%, **73.89%**, 6e<sup>-90</sup> | NADH10 |
|  | *orf198* | 24672-25268 | - | - | - |
|  | *rpl2* | 25343-26119 | *Cryptocaryon irritans* (YP010632200) | 95%, 36.11%, 7e<sup>-23</sup> | Ribosomal protein |
|  | *rps19* | 26139-26441 | - | - | Ribosomal protein |
|  | *rps13* | 26453-27424 | *Tetrahymena rostrata* (QBI37874) | 68%, 38.76%, 3e<sup>-30</sup> | Ribosomal protein |
|  | *orf171* | 28449-28964 | - | - | - |
|  | *nad9* | 28984-29610 | - | - | - |
|  | *rps12* | 29954-30349 | Protozoan of *Idotea baltica* (MCL4137936) | 75%, **66.67%**, 2e<sup>-37</sup> | Ribosomal protein |
|  | *orf169* | 30374-30883 | - | - | - |
|  | *cox2* | 30978-34934 | *Paramecium caudatum* (YP003734428) | 29%, 43.62%, 6e<sup>-37</sup> | COX2 |
|  | *orf265* | 35066-35863 | - | - | - |
|  | *orf474* | 37121-38545 | - | - | - |
| 2 | *orf699* | 4260-2161 | - | - | - |
|  | *orf525* | 10199-8622 | - | - | - |
|  | *orf549* | 11884-10235 | - | - | - |
|  | *atp-9* | 13388-13158 | *Tetrahymena paravorax* (YP740778) | 89%, **67.65%**, 1e<sup>-24</sup> | ATP-9 |

Data available from across the Ciliophora provided 23 potential comparators, which together included four directly comparable proteins (ATP9-COB-COX2-COX1) and were prepared into a maximum-likelihood phylogenetic tree (Fig. 5). The tree revealed that the novel species groups most closely with *M. luciensis* isolates MH182621 and AWT08281 (Fig. 5). Within this same group, we determined that a mitochondrial genome available from the host *Idotea baltica* (Isopoda; JAACYD010007607) also showed similarity and phylogenetic positioning, highlighting another putative ciliate of this group that may have gone undiagnosed.

**Fig 5.**
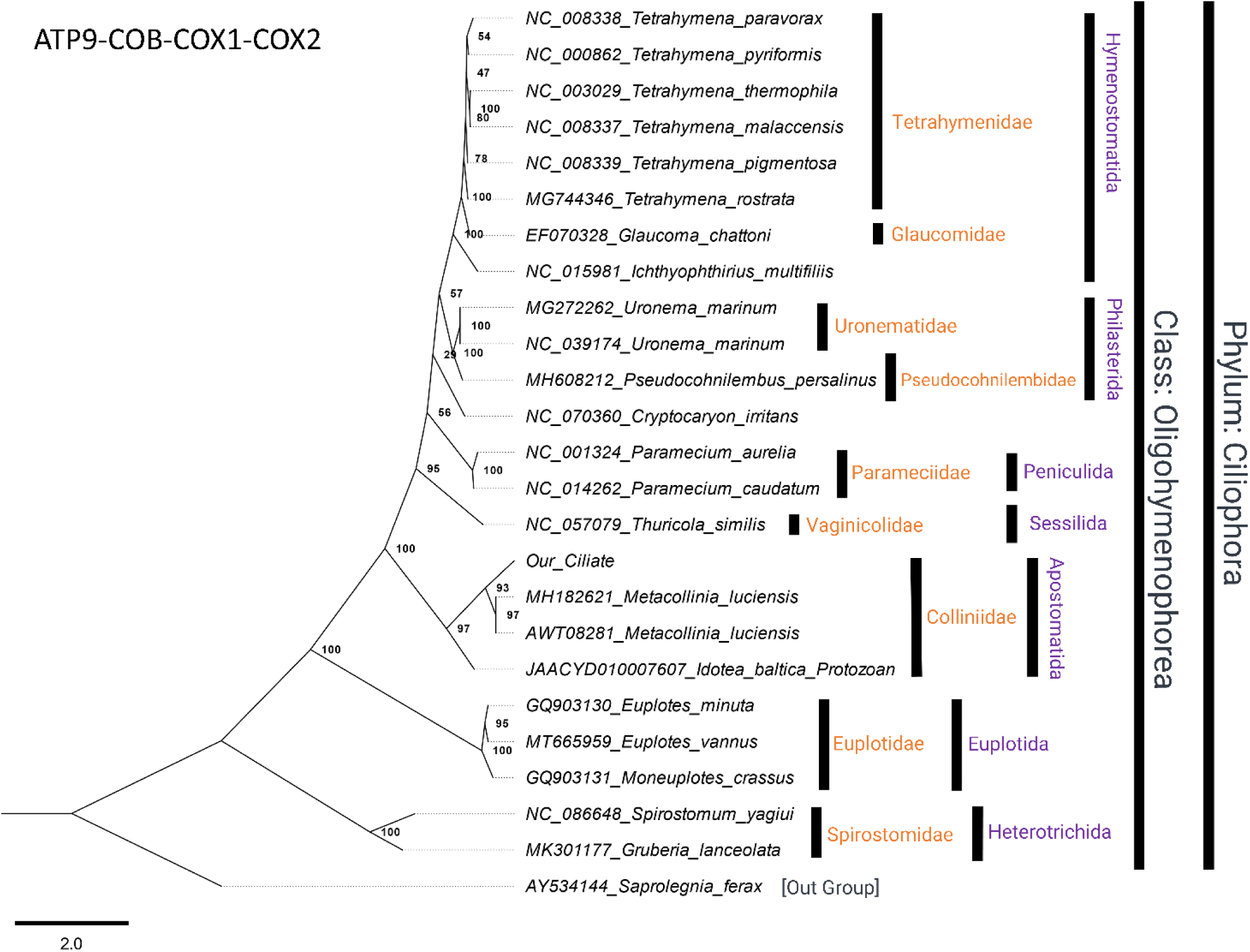
A maximum-likelihood tree based on four proteins (ATP9, COB, COX1, COX2), using mitochondrial sequences from across the Ciliophora. The tree includes 25 sequences (*Saprolegnia* is used as an outgroup) and includes 3509 columns, 2357 distinct patterns, 1548 parsimony-informative, 595 singleton sites, 1366 constant sites. Best-fit model: mtZOA+F+I+G4 chosen according to Bayesian Information Criterion. Log-likelihood of consensus tree: -46022.229.

Nuclear genes were not assessed from the metagenomic data during this study; however, information was gathered from the data to support the PCR diagnostic used to reproduce the 18S sequence for this species, as described above.

## 4. Taxonomic summary

Kingdom: Protozoa

Superphylum: Alveolata

Phylum: Ciliophora Doflein, 1901

Subphylum: Intramacronucleata Lynn, 1996

Class: Oligohymenophorea de Puytorac et al. (1974) Order: Apostomatida Chatton and Lwoff, 1928 Family: Colliniidae Cepede, 1910

Species: *Metacollinia emscheri* n. sp. Prati, Rückert, Grabner, Sures, Bojko, 2024

Formal taxonomic summary: The tomonts are ∼67 µm long with 40-47 (possibly more) slightly spiraled kineties, a non-ciliated cortical band, and 6-10 contractile vacuoles arranged either in two parallel rows or scattered across the cytoplasm. At the end of palintomy, during which the number of kineties decreases to nine, oral fragments (buccal kineties x, y, and z) appear.

Type locality: Boye catchment, North-Rhine Westphalia, Germany (Boye 51.553107; 6.948059 and 51.534515; 6.997740, Haarbach 51.570306; 6.960891 and 51.562638; 6.955581)

Type host: *Gammarus pulex* Linnaeus, 1758 (Crustacea, Malacostraca, Amphipoda, Gammaridae) Type habitat: Endoparasitic, found in the hemolymph and muscle tissues of hosts collected in lowland streams at the type locality.

Holotype: The name-bearing type of this species is the specimen illustrated in Fig. 2 and 3 (ICZN 1999, Articles 73.1.4).

DNA sequences: 18S rRNA and the draft mitogenome are publicly available via GenBank (18S PQ3206 - PQ3208, mitogenome: .............)

Etymology: The species epithet emscheri refers to the Emscher River catchment, which comprises the type localities found in its tributary, the Boye and the Haarbach.

## 5. Discussion

This study describes a novel sanguicolous apostome ciliate that infects the amphipods *G. pulex* and *G. fossarum*, from four populations of the Boye (North Rhine-Westphalia, Germany). Our phylogenetic analyses place *M*. *emscheri* n. sp. within the Collinidae family. Due to morphological characteristics, phylogenetic similarity with the type species, *M*. *lucensis*, and the absence of DNA sequences of less molecularly divergent species, we provisionally place this ciliate in the genus *Metacollinia*. This concept is supported by *18s*, *cox1,* and a concatenated multi-protein phylogenetic approach (Fig. 4-5).

### 5.1 Prevalence of *Metacollinia emscheri* n. sp

The presence of *M. emscheri* n. sp. in *G. pulex* clade E, *G. pulex* clade C, and *G. fossarum* clade C2 indicate lower host specificity than other collinids, as these have mostly been reported from single hosts. The prevalence of *M. emscheri* n. sp. across the four infected amphipod populations indicates medium infection levels and, in the case of the population collected from the lower stretch of the Haarbach (HAAun) temporal stability. This sanguicolous ciliate appears to be localized in the Haarbach and the lower part of the Boye, since none of the amphipods collected upstream of BOYohKI (located just above the junction of the two streams) were infected. However, we cannot be certain that *M. emscheri* n. sp. is absent from the upper stretch of the Boye catchment. This ciliate might be present in lower abundances and have evaded detection, due to a relatively small sample size (∼ 30 individuals). It is plausible that seasonal dynamics may also play a role - for example, the prevalence of *P. branchiarum* in *G. pulex* collected in a locality in proximity to Prague varied between 1% in January and 20% in August (De Puytorac and Lom, 1962). The close proximity of BOYuhHA, the most downstream site, with the River Emscher (2.7 km), suggests that *M. emscheri* n. sp. might also be present in the latter.

Assuming that the observed replacement of the muscle mass in infected amphipods may hinder the host’s ability to withstand water flow, it is plausible that a good portion of infected individuals passively drift downstream and end up in the River Emscher. Furthermore, all infected amphipods had relatively large sizes, which alone might influence amphipod drift. Accordingly, both large-sized and microsporidian-infected *G. pulex* in a nearby watercourse, the Rotbach, drifted more than smaller and uninfected individuals (Prati et al., 2023). The exclusive presence of *M*. *emscheri* n. sp. in relatively large-sized amphipods is not surprising. Generally, parasite prevalence in aquatic organisms, including amphipods, is positively correlated with host size, as older individuals are more likely to be exposed to infections over their lifespan (Poulin, 2000; Prati et al., 2023; Soldánová et al., 2010). Furthermore, size-asymmetric cannibalism, which is common in many aquatic taxa, including *G. pulex*, may potentially favor the transmission of parasites via ingestion of infected individuals (Arundell et al., 2019a, 2019b; MacNeil et al., 2003; McGrath et al., 2007).

### 5.2 Transmission of *Metacollinia emscheri* n. sp

The transmission of *M*. *emscheri* n. sp., like that of other members of the Collinidae and Pseudocollinidae families, remains unknown (Chantangsi et al., 2013; Gómez-Gutiérrez et al., 2012; Lynn, 2008; Metz and Hechinger, 2021). Poisson (1921) described *M. luciensis* from the digestive tract of *Orchestia gammarellus*, but mentioned that the ciliate might be able to pierce through the intestine wall and penetrate the coelom. Balbiani (in Cépède, 1910) hypothesized that host injury is required for ciliate transmission. Such a pathway might also be possible for *G. pulex* as injuries often occur in adult individuals, particularly during mating, when males compete to assert their dominance over females (Dick and Elwood, 1990).

Alternatively, pre-existing parasitic infections might favor secondary infections by hindering the host immune system (Cox, 2001). This does not seem to be the case in the present study since most infected amphipods did not show co-infections; similarly, Bojko et al. (2017) noted a few additional parasites present alongside their ciliate in *G. roeselii*, which was sampled in Poland and observed at a prevalence of <1%. A more plausible theory has been proposed by Metz and Hechinger (2021), who, after failing multiple attempts to experimentally infect healthy and injured crabs with *L*. *grapsolytica* (via exposure to swimming tomites, infected conspecifics, or direct injection of life cells) proposed that the parasite has a complex life cycle with development outside the host. This might possibly involve an intermediate host. Such a possibility might also apply to *M. emscheri* n. sp.; however, more extensive studies are required to shed light on its life cycle. Given that we have identified proteins from the mitochondrial genome with similarity to a protozoan symbiont of *Idotea baltica*, it may be that isopods can also host this group of parasitic ciliates.

### 5.3 Ecological importance of *Metacollinia emscheri* n. sp

Irrespective of its elusive life cycle, *M. emscheri* n. sp. is of high ecological importance. With prevalences up to 21% in the infected populations, this parasite is likely to strongly affect host population dynamics. Due to reduced muscular mass, host movement and activities such as feeding and mating might be affected in individuals with high infection levels. As amphipods are keystone species in freshwater aquatic environments, *M. emscheri* n. sp. may indirectly impact ecosystem functioning. For instance, infected individuals may experience reduced shredding activity and, consequently, a slowdown of leaf-litter breakdown decreases the amount of nutrients and processed material available for microorganisms and other invertebrates. Such a slowdown might be particularly detrimental to small stream food webs as the decomposition of terrestrial organic matter often constitutes the main source of energy (Gessner et al., 1999; Wallace et al., 1997). Furthermore, if *M. emscheri* n. sp. directly or indirectly increases the mortality rate in amphipods, species that rely on such prey may face reduced food availability. Gammarus spp. are an especially important food source for fish and, unlike other macroinvertebrates, are available year-round (Macneil et al., 1999).

### 5.4 Taxonomy and nomenclature of the family Collinidae

Jankowski (2007) undertook a comprehensive revision of the nomenclature and taxonomy of Collinidae, suggesting that the genus *Collinia* should be split into three distinct genera: *Collinia*, *Paracollinia*, and *Metacollinia*. *Collinia* (type species: *Collinia circulans*) was originally observed by Balbiani 1885 (in Cépède, 1910) in the freshwater isopod, *Asellus aquaticus*. *Collinia circulans* is characterized by a large tomont (70-80 µm long and 30-35 µm wide) with ∼ 13 spiraled bipolar kineties and a single row of ∼ 12 contractile vacuoles. The tomite is small, showing short x, y, and z kineties reminiscent of those of the foettingeriids (De Puytorac and Lom, 1962; Lynn, 2008; Lynn and Strüder-Kypke, 2019).

The genus *Paracollinia* (type species: *Paracollinia branchiarum*) was observed in 1852 by Stein, to infect the freshwater amphipod *G. pulex*. It has large tomonts with 55 to 60 straight evenly spaced kineties without a non-ciliated cortical band and a smaller flattened tomite with 9 kineties and short x, y, and z kineties, reminiscent of those of the foettingeriids (Jankowski, 2007; Lynn and Strüder-Kypke, 2019).

The genus *Metacollinia* (type species: *Metacollinia luciensis*) was observed in the marine amphipod *O. gammarellus* by Poisson (1921). It has large tomonts with 65 to 70 slightly spiraling kineties, a large non-ciliated cortical band and two lateral rows of 8-10 contractile vacuoles. Like the previous species, the tomite has 9 kineties and short x, y, and z kineties, reminiscent of those of the foettingeriids (Jankowski, 2007; Lynn and Strüder-Kypke, 2019).

More recently, a novel genus, *Lynn*i*a*, (type species: *L*. *Grapsolytica*) found in the marine crab *Pachygrapsus crassipes* has been included in the Collinidae (Metz and Hechinger, 2021). The tomont is large with ∼ 11 equidistant spiraled bipolar kineties, lacks a non-ciliated cortical band, and possesses randomly scattered subpellicular contractile vacuoles. The tomite is small, showing short x, y, and z kineties reminiscent of those of the foettingeriids; however, unique compared to any known apostome, the eighth and ninth kineties are both interrupted, and once impregnated with silver, the tomite presents a hook-shaped arrangement of ventral somatic kineties (Metz and Hechinger, 2021).

Compared to *L*. *grapsolytica*, the tomite of *M*. *emscheri* n. sp. bears more resemblance with that of *C. ciruclans*, *P. branchiarum,* and *M*. *luciensis*. Only the ninth somatic kinety in *M. emscheri* n. sp. is interrupted, and no hook-shaped arrangement of ventral somatic kineties is present. Like *M*. *luciensis*, the tomont of *M*. *emscheri* n. sp. displayed slightly spiraled kineties and a non-ciliated cortical band, suggesting the placement of the ciliate in this genus. The average length of tomont in *M*. *emscheri* n. sp. (∼67 µm) was similar to that reported in *M*. *luciensis* (∼70 µm) by De Puytorac and Grain (1975). However, the kineties, numbering 40-47, were lower than the 65-70 reported in *M*. *luciensis.* It is possible that the smaller size and lower number of kineties observed in *M*. *emscheri* n. sp. might be due to regression processes occurring during the tomitogenesis, as reported for *M*. *luciensis* collected by Lynn and Strüder-Kypke (2019). Interestingly, the contractile vacuoles in the tomont of *M*. *emscheri* n. sp., which numbers 6-10 in contrast to 8-10 found in *M*. *luciensis* (De Puytorac and Grain, 1975), are arranged either in two parallel rows or scattered across the cytoplasm.

Phylogenetic analyses supports these morphological deductions, and indicates that *M*. *emscheri* n. sp. is more closely related to *M*. *lucensis* than *L*. *grapsolytica*, the only other Collinidae for which DNA sequences are available. In the absence of DNA sequences of species belonging to the genera *Collinia* and *Paracollinia*, and based on morphological similarity with *M*. *lucensis*, we provisionally place *M*. *emscheri* n. sp. in the genus *Metacollina*. Given the gap in DNA sequence reference data, we cannot exclude that the proposed placement might change once more sequences of Collinidae species become available.

## Authors contribution

Sebastian Prati, Daniel S. Grabner, Bernd Sures, and Jamie Bojko conceived the study. Sebastian Prati carried out the sampling. Sebastian Prati, Sonja Rückert, and Jamie Bojko performed laboratory analyses. Jamie Bojko conducted the metagenomic and bioinformatic analysis and supervised the study. Sebastian Prati wrote the first draft of the manuscript, and all authors contributed critically to the final draft. All authors read and approved the final manuscript.

## Acknowledgments

This study was performed within the Collaborative Research Center (CRC) RESIST (A09) and funded by the Deutsche Forschungsgemeinschaft (DFG, German Research Foundation) – CRC 1439/1 – project number: 426547801. We are grateful to the Developmental Biology group of the University of Duisburg-Essen and, in particular, to Sabine Schneider for providing access and assistance with the histological instruments. We thank the Imaging Center Essen (IMCES) at the Faculty of Medicine of the University of Duisburg-Essen, in particular Bernd Walkenfortfor for providing access and assistance with the SEM equipment.

## Appendix A. Supplementary data

Supplementary material 1

