## Supplementary material 1 for "*Metacollinia emscheri* n. sp., a novel sanguicolous apostome ciliate of freshwater amphipods (*Gammarus* spp.)"

**Genetic and morphological characterization of *Metacollinia emscheri* n. sp., a novel sanguicolous apostome ciliate of the freshwater amphipods *Gammarus pulex* and *G. fossarum***

Sebastian Prati[1,2]*, Sonja Rückert[2,3], Daniel S. Grabner[1,2], Bernd Sures[1,2,4], Jamie Bojko[5,6]

[1]Aquatic Ecology, University of Duisburg-Essen, Universitaetsstr. 5, 45141 Essen, Germany. [2]Centre for Water and Environmental Research, University of Duisburg-Essen, Universitaetsstr. 5, 45141 Essen, Germany. [3]Eukaryotic Microbiology, University of Duisburg-Essen, Universitaetsstr. 5, 45141 Essen, Germany. [4]Research Center One Health Ruhr of the University Alliance Ruhr, University of Duisburg-Essen, Universitätsstraße 5, 45141 Essen, Germany. [5]National Horizons Centre, Teesside University, Darlington, DL1 1HG, United Kingdom. [6]School of Health and Life Sciences, Teesside University, Middlesbrough, TS1 3BX, United Kingdom

### Quantitative Protargol Staining protocol (version 10/02/2024)

**Solutions needed:**

1% agarose: 0.1g in 10 ml MilliQ. Prepare at each run.

10% formaldeyde: 2.7 ml formaldeyde 37%, top up with MilliQ water to 10 ml. Prepare at each run.

0.2% potassium permanganate: 0.02g in 10 ml MilliQ. Prepare at each run.

2.5% oxalic acid: 0.25g in 10 ml Millq. Prepare at each run.

Silver proteinate (~1% protargol): prepare a 0.3% sodium hydroxide solution: 0.03g NaOH in 10 ml MilliQ and a 1.26% silver nitrate solution: 0.126g silver nitrate in 10 MilliQ. These solutions can be stored in the fridge. To prepare the silver proteinate solution, sprinkle 0.085 g of powdered proteose peptone on the surface of 8 ml MilliQ water and let it dissolve without stirring. Add 1 ml of 0.3% sodium hydroxide (NaOH) solution and mix thoroughly, then add dropwise 1 ml of a 1.26% solution of silver nitrate. Test the pH, and adjust to 8 with 0.3% sodium hydroxide solution (0.4-1 ml). Prepare at each run

Developer: 0.17 g sodium sulfite (Na2SO3) in 10 MilliQ, add 0.01g of hydroquinone

5% sodium thiosulfate: 0.50 g sodium thiosulfate $Na_2S_2O_3 \cdot 5H_2O$ in 20 ml MilliQ. Prepare at each run.

**Protocol:**

1) Isolate the ciliates with a glass pipette
2) Rinse the fixative by transferring the samples from one convex microscope glass containing distilled water to another 4-5 times
3) Place the ciliates in the middle of a 25mm cellulose nitrate filter 0.45 micron held on a filter holder connected to a syringe
4) Using the syringe, suck gently all the water from the filter, but do not overdue, or the cell may get ruined
5) Remove the filter and cut the excesses. Quickly place a drop of 1% liquid agarose (kept at 60°C) on a warm microscope glass (40°C) and place the filter on it. Add a drop of 1% liquid agarose on a cover slip and quickly put it over the filter, applying gentle pressure to obtain a thin and even layer of agarose
6) Allow the agarose to solidify (15 min)
7) Remove excess agarose and transfer the filter into 10% formaldehyde for 10 min to increase the melting point of agarose
8) Transfer in a 0.2% solution of potassium permanganate for 5 minutes for bleaching
9) Wash in distilled water by swirling the filter for 30 sec to remove the excess potassium permanganate
10) Place the filter in 2.5% oxalic acid for 5 min to completely remove potassium permanganate, which inhibits staining
11) Wash in distilled water for 10 min to remove the oxalic acid, which inhibits staining
12) Place filters in the silver proteinate solution heated at 60°C for 30-40 minutes, control the color. If not strong enough, let it for longer, then let it cool down for 10 minutes to room temperature
13) Transfer the filter to the developer solution for 10-40 sec until a brown color emerges on target structures. The concentration of the developer can control the intensity of the stain.
14) Transfer the filter into distilled water and check the coloration of the target structures to evaluate the staining. If the structures are insufficiently stained, go back to point 11; if they are overly stained, go back to point 9; otherwise, continue with the next step.
15) Place filters in 5% sodium thiosulfate solution for 3 min to fix the stain, not that the color will darken a little
16) Wash in tap water for 10 min
17) Dehydrate in isopropyl alcohol series (not ethanol to reduce filter´s warping) (50-70-90-95-100-100%) for 5 min each
18) Transfer to 50% Rothiclear and 50% isopropyl alcohol for 5 min
19) Transfer 100% Rothiclear for 5 min twice. This makes the filters transparent and clears the cells.
20) Mount each filter on a slide by placing drops of Histokitt II below the filter and on the coverslip. Ensure that no air is trapped under the filter or coverslip. Apply light pressure to remove excessive mounting media. Let it dry.
